# The *SINGLE FLOWER (SFL)* gene encodes a MYB transcription factor that regulates the number of flowers produced by the inflorescence of chickpea

**DOI:** 10.1101/2020.12.19.423578

**Authors:** Cristina Caballo, Ana Berbel, Raul Ortega, Juan Gil, Teresa Millán, Josefa Rubio, Francisco Madueño

## Abstract

**research conducted & rationale:** Legume species usually have compound inflorescences, where flowers appear in secondary inflorescences (I2), at lateral positions of the primary inflorescence (I1), in contrast to simple inflorescences, as in *Arabidopsis*, where flowers are formed in the primary inflorescence stem. The number of flowers per I2, characteristic of each legume species, determines inflorescence diversity, and the number of pods produced, which can affect yield. Gene Regulatory Network that controls the activity of I2 meristems, and therefore the number of flowers per secondary inflorescence is mostly unknown, as well as how specific are factors controlling this trait and whether they share this function in other meristems.

**methods:** Chickpea produces one flower per I2 but *single flower* (*sfl*) mutants produce two (double-pod phenotype). By mapping the *sfl-d* mutation and identification and analysis of a second mutant allele we have isolated *SFL*. We used scanning electron microscopy to study the effect of *sfl* mutations on inflorescence ontogeny and *in situ* hybridization to study the expression of *SFL* and of meristem identity genes in the developing chickpea inflorescence.

**key result:** We show that the *SFL* gene corresponds to *CaRAX1/2a*, encoding a MYB transcription factor. Our results show that *CaRAX1/2a / SFL* is specifically expressed in the I2 meristem, possibly activated by *CaVEGETATIVE1*.

**main conclusion & key points for discussion:** Our findings reveal that *SFL* plays a central role in the control of chickpea inflorescence architecture, specifically acting in the I2 meristem to control the time length for which it is active, and therefore determining the number of floral meristems that it can produce.

## INTRODUCTION

Inflorescence architecture is a relevant question in plant developmental biology, as it shapes plant form (Weberling, 1992; Benlloch et al., 2007). Moreover, inflorescence architecture is also relevant in agriculture, because it strongly influences fruit and seed production (Wang and Li 2008). Architecture of the inflorescence depends on the identity of the meristems in the inflorescence apex, which determines in which positions in the inflorescence axes flowers are formed, and on the activity of the inflorescence meristems, which control how many flowers are produced (Benlloch et al., 2007; Prusinkiewick et al., 2007; Teo et al., 2014).

While in simple inflorescences, such as that of *Arabidopsis*, flowers are directly formed in the primary inflorescence (I1) stem, the basic type of inflorescence in legumes is the compound inflorescence, where flowers (and therefore pods) appear in secondary inflorescence (I2) stems, formed in lateral positions of the primary inflorescence (Fig. **1a**,**b**; **Fig. S**; Benlloch et al., 2015). The development of the legume compound inflorescence depends on the I2 meristem identity gene *VEGETATIVE1* (*VEG1*), which encodes a MADS transcription factor without a clear functional homologue in Arabidopsis (Berbel et al., 2012; Cheng et al., 2018), and on the floral identity gene *PROLIFERATING INFLORESCENCE MERISTEM* (*PIM*), encoding a homologue of the Arabidopsis MADS transcription factor APETALA1 (AP1) (Mandel et al., 1992; Berbel et al, 2001; Taylor et al., 2002; Benlloch et al., 2006). The legume I2 meristems are, generally short-lived meristems that grow producing a number of floral meristems before degenerating into a residual organ or stub (**Fig. 1e**).

**Fig 1.**
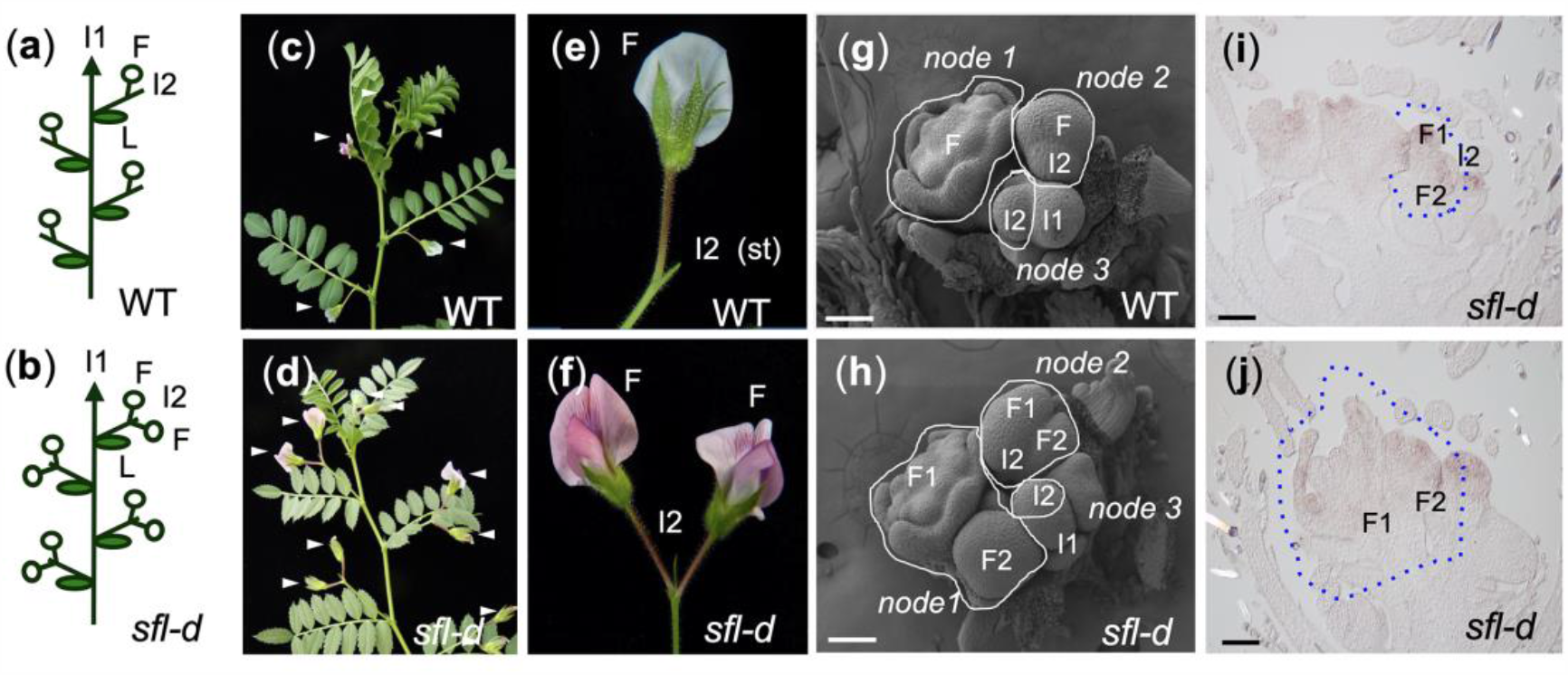
Double-pod phenotype in chickpea caused by mutation in the *SINGLE FLOWER* (*SFL*) gene and ontogeny of the inflorescence of the *sfl-d* mutant. (a, b) Diagrams of wild-type (WT) and *sfl-d* chickpea plants. Flowers (F) develop at secondary inflorescences (I2) that are formed in the axil of the leaves (L) of the primary inflorescence (I1) stem. Wild-type I2s (a) produce one flower while *slf-d* I2s (b) produce two flowers. (c, d) Wild-type and *sfl-d* chickpea plants. Arrowheads mark individual flowers formed at the I2s of the wild-type (c) and two flowers in the *sfl-d* (d) I2s. (e) Close-up of a wild-type I2, where the stub (st) is marked. (f) Close-up of a *slf-d* I2. (g) Scanning electron micrograph (SEM) of the inflorescence apex of a wild-type plant. In each I2 node one flower is found. (h) SEM of the inflorescence apex of a *slf-d* plant. In the I2 nodes two flowers (at different developmental stages) are found. (i, j) *In situ* hybridization of *CaPIM* mRNA in inflorescence apices of the *sfl-d* mutant, where each I2 node bear two flowers at different developmental stages.

**Fig 2.**
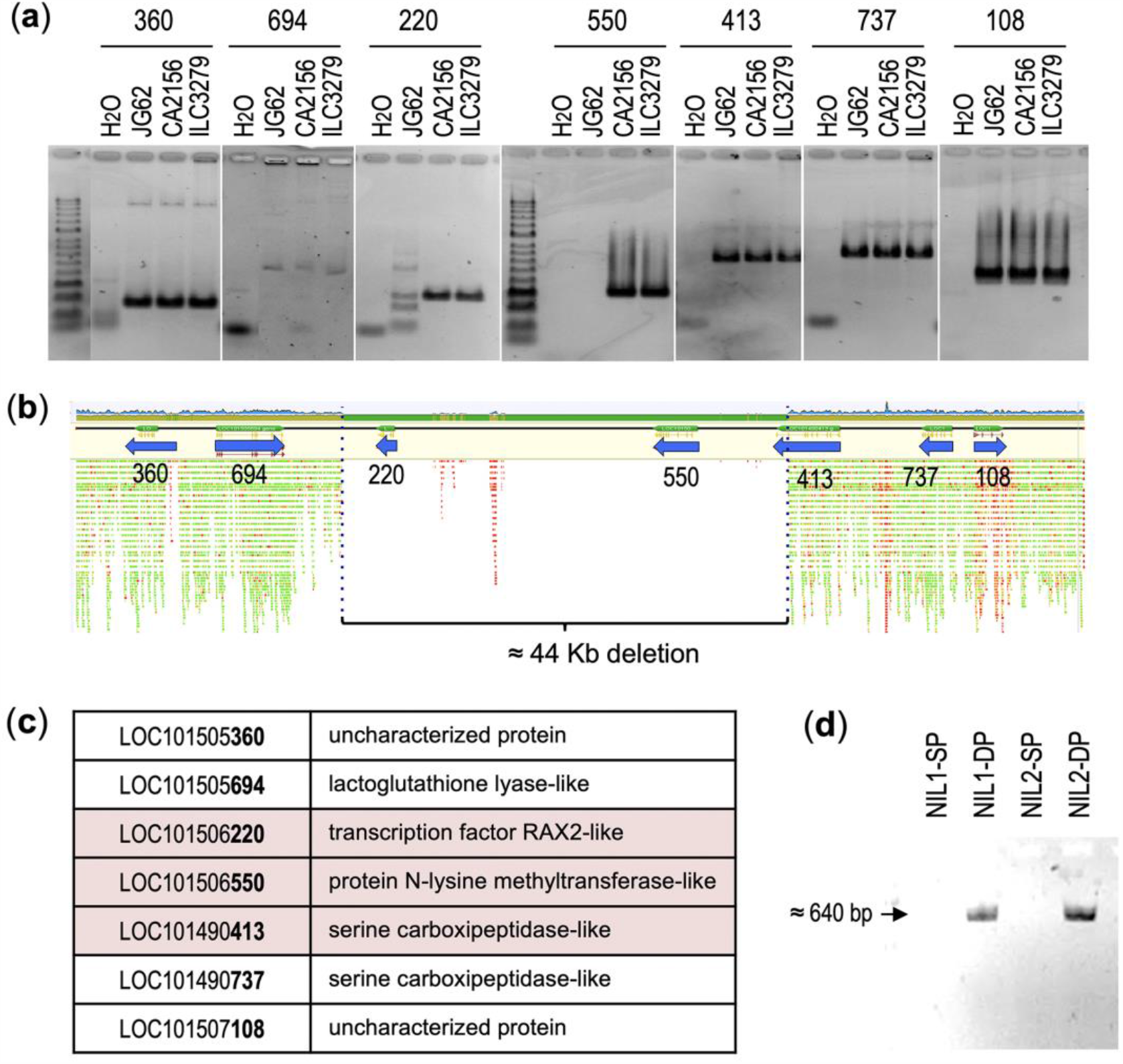
Deletion in the 92.6 kb *SFL* mapping interval. (a) PCR amplification with primers for the seven genes in the 92.6 kb *SFL* mapping in DNA from the double-pod line JG62 and the single-pod lines CA2156 and ILC327. (b) Mapping of sequencing reads of the JG62 re-sequencing against the genome of the reference single-pod line CDC-Frontier in the 92.6 kb *SFL* mapping interval, showing a deletion affecting three genes. (c) List of the genes contained in the 92.6 kb *SFL* mapping interval. Genes affected by the deletion are highlighted in pink. (d) PCR amplification with primers at the limit of the deletion in two pairs of single-pod (SP) or double-pod (DP) nearly isogenic lines (NIL1, NIL2). Scale bars: 100μm.

**Fig 3.**
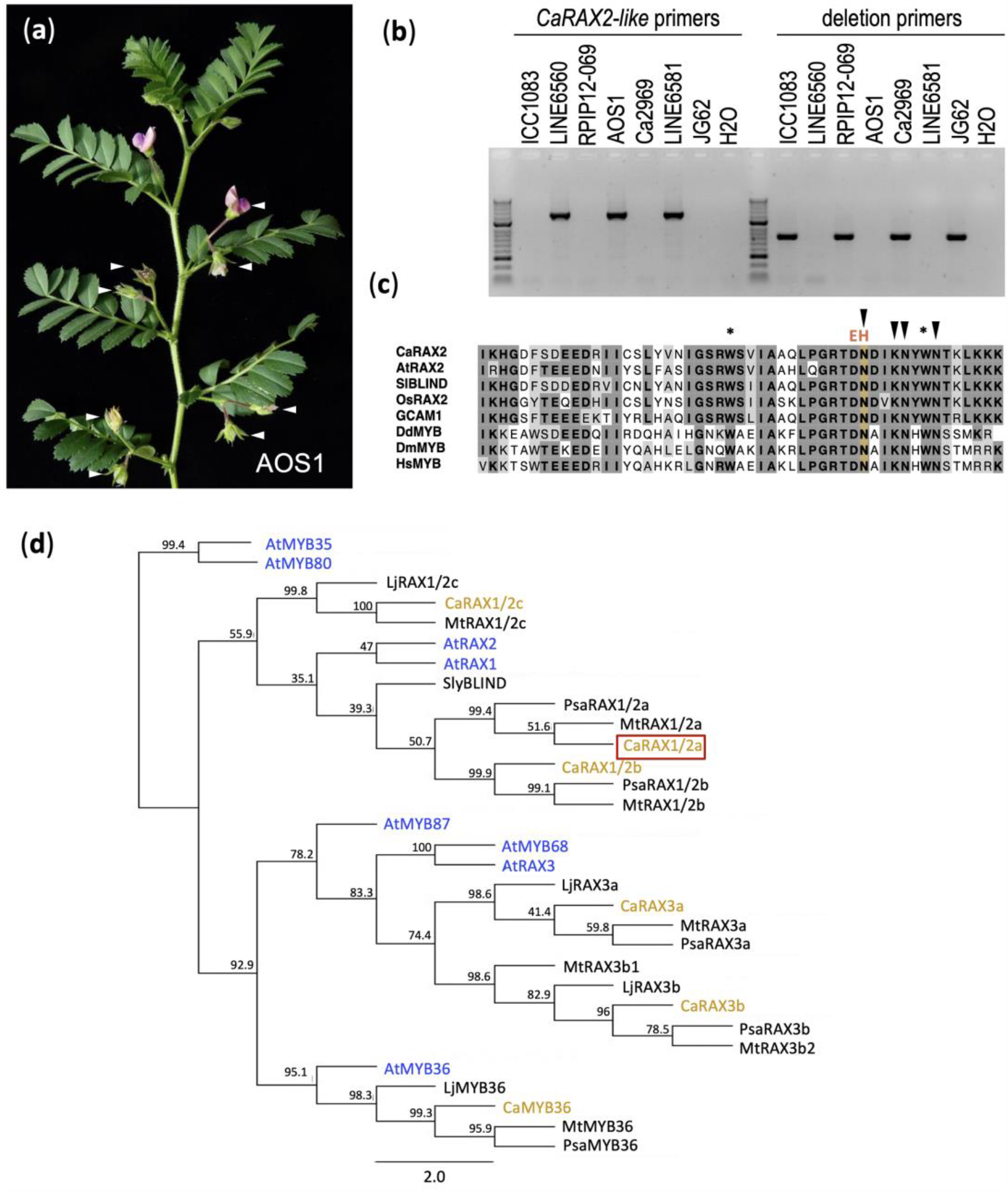
*CaRAX2-like* is mutated in a new mutant allele of *SFL* and phylogenetic tree of legume RAX proteins. (a) Double-pod phenotype of an AOS1 plant. Arrows mark the flowers. (b) PCR amplification in the USDA double-pod lines with primers pairs at *CaRAX2-like* or at the limits of the deletion. (c). ClustalW alignment of the R3 repeat of representative R2R3 MYB proteins from plants, microorganisms, and animals. Ca, Cicer arietinum; At, *Arabidopsis thaliana;* Sl, *Solanum licopersicum*; Os, *Oriza sativa*; GCMA1, *Marchantia polimorpha*; Dd, *Dictioustelium discoideum*; Dm, *Drosophila melanogaster*; Hs, *Homo sapiens*. Asterisk mark conserved triptophan residues. Arrowheads mark base-contacting residues of the mouse homologue of Hs MYB (d) Phylogenetic tree of subgroup 14 MYB proteins from Arabidopsis and legumes. Legume proteins have been named after their Arabidopsis Homologues. CaRAX2-like / CaRAX1/2a is framed in red. At, *Arabidopsis thaliana*; Ca, *Cicer arietinum* (chikpea); Sl, *Solanum licopersicum*; Lj, *Lotus japonicus*; Psa, *Pisum sativum* (pea); Mt, *Medicago truncatula*. Accession number of genes in the phylogenetic tree can be found in Table **S2**.

**Fig 4.**
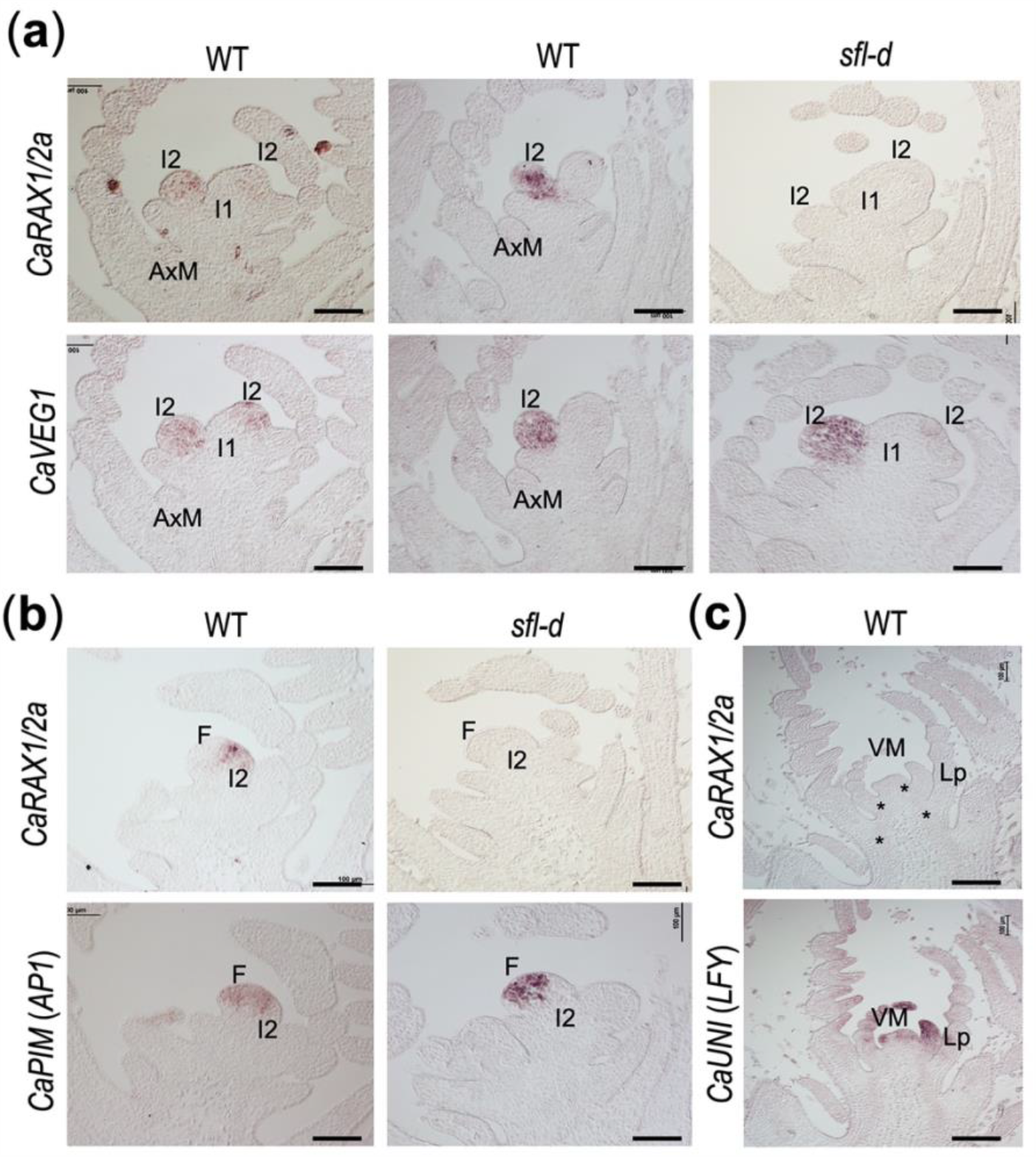
Expression pattern of the *CaRAX1/2a / SFL* gene in shoot apices of chickpea. (a) *In situ hybridization* of *CaRAX1/2a* mRNA (top row) and of the secondary inflorescence meristem (I2) marker *CaVEG1* mRNA (lower row) in contiguous sections of inflorescence shoot apices of wild-type (WT) or *sfl-d* plants. I1, primary inflorescence meristem; AxM, axillary meristem. (b) *In situ hybridization* of *CaRAX1/2a* mRNA (top row) and of the floral meristem (F) marker *CaPIM* mRNA (lower row) in contiguous sections of inflorescence shoot apices of wild-type or *sfl-d* plants. (c) *In situ* hybridization of *CaRAX1/2a* mRNA (top) and of the leaf and floral marker *CaUNI* mRNA (bottom) in contiguous sections of vegetative shoot apices of a wild-type plant. Asterisks in the top image mark leaf axils. VM, vegetative shoot apical meristem; Lp, leaf primordia. Scale bars: 100μm.

The number of flowers formed by an inflorescence meristem can depend on its size and division pattern (Clark et al., 1995), or on the length of time for which it stays active, producing floral meristems (Balanzá et al. 2018). The number of flowers per I2 is a relevant trait for at least two reasons. First, it is a key factor that creates diversity in inflorescence architecture and shape. Thus, the number of flowers per I2 is characteristic of each legume species and variety. For example, while *Pisum sativum* (pea) and *Medicago truncatula* I2s form 1-2 flowers, *Medicago sativa* (alfalfa) I2s produce 8-12 flowers and *Vicia cracca* (cow vetch) I2s produce dozens of flowers (Fig. **S1;** Benlloch et al., 2015). Second, the number of flowers per I2 determines the number of pods produced by the plant, having the potential to positively impact on yield (Kumar, 2000; Ali et al., 2010). Mutations whose only effect is to increase the number of flowers per I2 have been described in several legumes (Murfet, 1985; Srinivasan et al., 2006; Mishra et al., 2019), indicating the existence of genes specifically regulating this trait. In spite of the relevance of this trait, and while quite much is known about the genetic control of meristem identity in the legume inflorescence (Berbel et al, 2012; Benlloch, 2015; Cheng et al., 2018) the regulation of the number of flowers per I2 is poorly known and, until now, no gene that specifically controls I2 meristem activity has been isolated.

How is the number of flowers produced by the legume I2s controlled? In pea, two genes have been described, *Fn* and *Fna*, whose recessive mutations lead to an increase in the number of flowers/pods formed in the I2, resulting in the multipod trait (White, 1917; Lamprecht, 1947). Similarly, in *Cicer arietinum* (chickpea) two genes, *CYM* (*CYMOSE*) and *SFL* (*SINGLE FLOWER*) have been described whose recessive mutations specifically lead to the formation of a larger number of flowers by the I2s (Srinivasan et al., 2006).

In chickpea, while most genotypes produce just one flower per I2 (“wild-type”; Fig. **1a**,**c**,**e**) (Prenner, 2012), mutations in the *SFL* gene lead to plants whose only evident phenotype is the production of two flowers/pods per I2 (double-pod trait; Fig. **1b**,**d**,**f**), though, occasionally, three flowers per I2 can be observed in *sfl* mutants (Srinivasan et al., 2006). Some studies correlate the *sfl* double-pod mutation with a higher seed yield, while others with higher yield stability, supporting a positive effect of the double-pod mutation on crop performance and its agronomical potential (Sheldrake et al., 1978; Rubio et al., 1998; Kumar et al., 2000; Rubio et al., 2004). Thus, the *SFL* gene represents a tool potentially useful to improve yield but also to understand the regulation of I2 meristem activity and its contribution in the generation of morphological diversity in legumes.

Using recombinant inbred lines (RILs) and pairs of near isogenic lines (NILs) the *SFL* gene was recently fine-mapped to a region of 92.6 Kb region of chromosome 6, with seven annotated genes (Jain et al., 2013; Varshney et al., 2013; Ali et al., 2016). Here, we characterize the *sfl-d* mutant, analysing the ontogeny of its inflorescence and the expression of a floral meristem gene. Then, we identify *SFL* as a homologue to the Arabidopsis *RAX1/2* genes, encoding a R2R3 MYB transcription factor, and investigate how it functions by studying its expression in the chickpea inflorescence apex compared to other inflorescence meristem genes. Our findings reveal that the *SFL* gene plays a central role in the control of architecture of the chickpea inflorescence, specifically acting in the I2 meristem to control the time length for which it is active, and therefore determining the number of floral meristems that it can produce.

## MATERIALS AND METHODS

### Plant material

Different sources of double pod were used: (i) JG62 (syn. ICC 4951) is an Indian landrace maintained by International Crops Research for the Semi-Arid Tropics (ICRISAT). JG62 acted as parental in a cross with the single-pod line CA2156 to develop five pairs of nearly isogenic lines (NILs) that were used to map *SFL* (Ali et al., 2016). (ii) Six genotypes from United States Department of Agriculture (USDA), described as double-pod (https://npgsweb.ars-grin.gov/gringlobal/search.aspx), (AOS1, CA2969, ICC1083, LINE6560, LINE6581, RPIP12-069-06223) were used to look for a second *sfl* mutant allele.

Single-pod genotypes were kabuli type. CA2156 is a Spanish cultivar, ILC3279 a landrace from the former Soviet Union maintained by the International Center for Agricultural Research in the Dry Area (ICARDA) and BT6-17 is an advanced line, from our breeding program at IFAPA.

### Controlled crosses

Genetic crosses were performed as previously described in (Caballo et al. 2018).

### Genotyping and sequencing

DNA was isolated using the DNAeasy Plant Mini Kit according to the manufacturer’s instructions (Quiagen). PCR reactions were carried out using Phusion High-Fidelity DNA polymerase (Thermo Scientific™) with 20 ng of plant genomic DNA. PCR products were analysed using non-denaturing polyacrylamide. The primers used in this work are listed in Table **S1**

PCR amplification products were sequenced by Sanger technology, either directly after purification with SureClean (Bioline) or after cloning them in pGEMTeasy (Promega)

### Bioinformatics analysis

In order to find differences between sequences of double-pod and single-pod genotypes, re-sequencing of the double-podded accession JG62 was re-sequenced using Ion-Torrent technology, and was mapped against the chickpea reference genome (from the single podded cultivar CDC-Frontier) available at NCBI (Varshney et al. 2013). The Binary Alignment Map (BAM) files were inspected using Geneious® 8 software used to polymorphism in relevant regions of the genome between both lines.

To infer the extension of the deletion affecting Ca*RAX2-like*, all reads assembled to a region of the chickpea chromosome 6 spanning 100 kb and containing genes LOC101505360, LOC101505694, LOC101506220 (Ca*RAX2-like*), LOC101506550, LOC101490413, LOC101490737 and LOC101507108 were extracted and re-assembled using Geneious mapper and the following options: minimum mapping quality = 30; maximum gaps per read = 10%; maximum gap size = 50000; word length = 17 and Index word length = 14; maximum mismatches per read = 15%; maximum ambiguity = 2.

### Scanning electron microscopy (SEM)

For SEM, chickpea inflorescence apices were fixed in FAE (50% ethanol, 3.7% formaldehyde, 5% glacial acetic acid) at 4°C overnight in the dark. Samples were dehydrated with ethanol and critical point dried in liquid CO2 (Leica CPD300). Dried samples were sputter-coated with argon–platinum plasma at a distance of 6–7 cm and 45 mA intensity for 15 sec in a sputtering chamber (Leica Microsystems EM MED020). Scanning electron micrographs were acquired using an AURIGA compact FIB-SEM (Zeiss, http://www.zeiss.com/) at EHT = 1–2 kV.

### *In situ* hybridization

RNA *in situ* hybridization with digoxigenin-labelled probes was performed on 8-µm longitudinal paraffin sections of chickpea meristems as described (Ferrándiz et al., 2000). Ca*VEG1 CaPIM* and *CaUNI*, which share 86.4%, 96.2% and 87.9% amino acid identity with *VEG1, PIM* and *UNI* genes from pea (Hofer et al., 1997; Taylor et al, 2002, Berbel et al., 2012 ref), respectively, were retrieved from the chickpea genome database. RNA antisense and sense probes of *CaRAX2-like*, Ca*VEG1 CaPIM* and *CaUNI* were generated using as substrate a 346-bp fragment of *CaRAX2-like* (593–988 from ATG), a 460-bp fragment of *CaVEG1* (237–696 from ATG), a 764-bp fragment of *CaPIM* (208–971 from ATG) or a 1007-bp fragment of *CaUNI* (182–1188 from ATG) amplified by PCR and cloned into the pGEM-T Easy vector (Promega). Transcription of probes was carried using SP6 and T7 polymerases, respectively. Signal was detected as a purple precipitate when viewed under the light microscope.

## RESULTS

### Effect of the *sfl-d* mutation on chickpea development

As previously mentioned, the I2 of wild-type genotypes produce one flower before terminating in a stub (Fig. **1a**,**c**,**e**). However, in homozygous lines for the *sfl-d* mutation in the *SINGLE FLOWER* (*SFL*) gene, the I2s produce two flowers/pods, (Fig. **1b**,**d**,**f**). This double-pod phenotype is recessive and only plants homozygous for the *sfl-d* mutation produce two flowers in their I2s (Srinivasan et al; 2006).

The production of two flowers suggest that, in double-pod genotypes, either the I2 meristem is larger and divides producing more flowers, or that it is active for longer and it has the time to form more than one flower in a sequential manner. To address this question, we used scanning electron microscopy (SEM) to analyse the ontogeny of the inflorescences of the pair of nearly isogenic lines NIL5-1V and NIL5-2V, single- and double-pod, respectively, derived from a cross with a *sfl-d* parental, JG62 (Alí et al., 2016). In inflorescence apices of the single-pod plants, each inflorescence node showed an I2 meristem from which only a floral primordium had initiated (Fig. **1g**). In the inflorescence apex of the double-pod plants, the I2 meristems had a similar size than wild type but, in each I2 node, two flowers at a different stage of development were observed (Fig. **1h**)

In addition, *in situ* hybridization of sections of double-pod inflorescence apices probed with the floral marker *CaPIM* (chickpea homologue of the floral meristem identity genes *PIM* and *AP1*), showed the presence of two flowers at different developmental stages in the I2 nodes of the double-pod mutants (Fig. **1i**,**j)**, confirming the result of the SEM analysis.

These results show that the two floral meristems produced by the double-pod I2s are initiated sequentially and indicate that the I2 meristems of the double-pod/*sfl-d* mutant produce more flowers than the wild-type I2 meristems because they are active for longer and eventually degenerate forming a stub as in the wild type.

### Identification of candidates for the *SFL* gene

The *sfl-d* mutation had been previously mapped to a 92.6 kb region of chromosome 6, with 7 annotated genes, which code for 2 uncharacterized proteins, 4 enzymes and a transcription factor, *CaRAX2-like* (Fig. **2a**,**b**,**c**) (Ali et al. 2015). When primer pairs for the seven genes of the 92.6 kb region were tested by PCR in single-pod parental lines they amplified gene fragments of the expected size (Fig **2a**). In contrast, in the double-pod line JG62 no amplification of the LOC101506550 (*N-lysine methyltransferase-like*) gene was observed and the band pattern when trying to amplify the LOC101506220 (*CaRAX2-like*) gene was unclear (Fig **2a**).

*CaRAX2-like* encodes a R2R3 MYB transcription factor with sequence similarity to *REGULATOR OF AXILLARY MERISTEMS* (*RAX*) genes from Arabidopsis (Keller et al., 2006; Muller et al., 2006), and it was suggested by Ali et al. (2016) as a putative candidate for the *SFL* gene. Because we could not amplify that gene in the *sfl-d* double-pod parental line JG62, we tried to amplify the *CaRAX2-like* gene in different double- or single-pod lines, to assess whether it was or not present in double-pod genotypes. Amplification of *CaRAX2-like* resulted in a fragment with the expected size in all single-pod lines (≈ 900 bp) while in double-pod lines the fragment was smaller than predicted (≈ 700 bp). Sequencing of the two PCR fragments showed that, as expected, in all cases the amplicon derived from single-pod genotypes was identical to *CaRAX2-like*, while the PCR amplicon from double-pod genotypes was unspecific and matched Ca*RAX3-like* (LOC101491244). This supports that the *CaRAX2-like* gene is absent from the genome of the double-pod genotypes tested. As *CaRAX2-like* and N-lysine methyltransferase are contiguous genes, these results suggested the existence of a deletion affecting these genes in the 92.6 kb *SFL* mapping interval.

To explore this possibility further, we examined the mapping quality of the JG62 (*sfl-d*) re-sequencing against the reference genome in this particular region of chickpea chromosome 6. We found a region of around 44 kb with an unusually low density of reads, and those reads that mapped showed a low map quality score (Fig. **2b**). This observation strongly supports the presence of a large deletion in this region of chromosome 6. We then extracted all the reads mapped to a 100 kb region containing the *SFL* mapping interval (Ali et al., 2016), and performed a new mapping against the same reference but allowing a within-read gap of 50 kb, which allowed us to infer the deletion breakpoints (Fig. **b**). As expected, the predicted 44kb deletion includes the genes LOC101506220 (*CaRAX2-like*) and LOC101506550 (*N-lysine methyltransferase-like*); the deletion also included part of a third gene, LOC101490413 (*serine carboxypeptidase-like*) (Fig. **2b, c**). To validate the existence of the predicted 44kb deletion in the double-pod genotype JG62, we designed primers at the limits of that predicted deletion and tested them by PCR in genotypes differing in the single/double-pod character. As expected, due to the large size of the sequence to be amplified, no PCR product was obtained in single-pod genotypes. In contrast, in double-pod genotypes, with the same primers, a fragment of the expected size (642 bp), was amplified (Fig. **2d**), which was latter sequenced to confirm the deletion borders. Taken together, these results show the existence of a deletion affecting three genes in the mapping region of the *SFL* gene, only present in the double-pod genotypes, which points to these three genes as the most likely candidates for the double-pod phenotype in the *sfl-d* mutants.

### Identification and analysis of a new double-pod mutant allele confirms *CaRAX2-like* as the *SFL* gene

To assess whether any of the three genes affected by the deletion was in fact responsible for the double-pod phenotype of the *sfl-d* mutant lines, we looked for new *sfl* mutant alleles different to the *sfl-d* mutation present in JG62, searching for double-pod mutants that did not contain the 44 kb deletion. We analysed six chickpea accessions described as double-pod in the USDA collection (Fig. **3a**). Two primers pairs designed for the 44 kb deletion and for the *CaRAX2-like* gene were used to test if the USDA genotypes also had the deletion present in the parental *sfl-d* double-pod mutant line JG62. Three of the six double-pod USDA genotypes (ICC1083, RPIP12-069-06223 and CA2969) contained the 44 kb deletion (Fig. **3b**), indicating that in these genotypes the double-pod mutation was the same as in JG62. However, the other three double-pod genotypes, LINE6560, AOS1, and LINE6581, did not contain the deletion but showed amplification with the Ca*RAX2-like* primers (Fig. **3b**). This suggested that the double-pod phenotype of these three last genotypes could be due to mutations in the *SFL* gene different to that in JG62.

To determine the allelic relationship of the double-pod mutation in the USDA lines with the *sfl-d* double-pod mutation of JG62, we crossed JG62 with the USDA double-pod lines AOS1 and LINE6560. The F1 plants from these crosses exhibited a double-pod phenotype (Fig. **S3a**,**b**), showing that there is no complementation between these double-pod lines and the JG62 *sfl-d* double-pod mutant. Moreover, F1 plants derived from the cross between the double-pod line AOS1 x single-pod line BT6-17 exhibited a single-pod phenotype (Fig. **S3c**), confirming that the double-pod mutation in the AOS1 line is recessive, like the *sfl-d* mutation of JG62. As only the JG62 mutant contains the 44 kb deletion, these results indicate that the USDA lines AOS1 and LINE6560 bear mutation(s) in the *SFL* gene (responsible for the double-pod character) allelic to the *sfl-d* mutation in JG62; this new allele was named *sfld-3*.

The three genes affected by the 44 kb deletion, present in JG62 and other related double-pod genotypes, were sequenced in the USDA lines and subsequently aligned against the chickpea reference sequence. In the three USDA genotypes both LOC101506550 (*N-Lysine methyltransferase-like*) and LOC101490413 (*serine carboxipeptidase-like*) genes had a sequence identical to the reference, while the LOC101506220 (Ca*RAX2-like*) gene had a sequence variant, in its coding sequence, identical in the three double-pod lines. The Ca*RAX2-like* gene in the USDA lines showed a change in two bases compared to the reference, affecting bases 306 and 307 of the coding sequence, which varied from CA in the reference to AC in the double-pod lines. This change of bases produces a variation in the amino acids 102 and 103 of the coded protein, which change from aspartate-asparagine in the wild-type reference to glutamate-histidine in the USDA double-pod lines (Fig. **3c**).

Alignment of the sequence of CaRAX2-like from the double-pod USDA lines to those of other related R2R3 MYB proteins, from plants and other organisms, showed that the mutation in the protein from the double-pod USDA genotypes is located in the R3 repeat of the conserved MYB DNA-binding domain (Fig. **3c**, Dubos et al., 2010). The amino acid residues affected by the mutation in the CaRAX2-like from the double-pod USDA lines, aspartate-asparagine, are conserved in the MYB domain of plant RAX proteins but they are also conserved in many MYB proteins from other organisms, from yeast to humans (Fig. **3c**). Moreover, in the mouse homologue of the human c-MYB protein, this asparagine residue was shown to directly contact DNA (Ogata et al., 1994; Martin and Paz-Ares, 1997) and therefore is presumably critical for function, which strongly suggest that the mutation on the USDA double-pod lines significantly affects the activity the activity of the CaRAX2-like protein.

The fact that two independent allelic mutations in the gene for the transcription factor CaRAX2-like, deletion and likely loss-of-function, associate to the double-pod mutant phenotype strongly indicates that *CaRAX2-like* corresponds to the *SFL* gene, responsible for the double-pod trait.

Phylogenetic analysis confirmed that, in agreement with its annotation, CaRAX2-like belongs to the large family of R2R3 MYB transcription factors, grouping with the subgroup 14, where other Arabidopsis RAX and the tomato Blind proteins are included (Fig. **3d**; Schmitz et al., 2002; Dubos et al., 2010). CaRAX2-like belongs to the same clade as AtRAX1 and AtRAX2, from Arabidopsis, although it groups together with a separate clade specific for legumes. As CaRAX2-like seems similarly close to both Arabidopsis proteins, and there is another CaRAX protein similarly close to AtRAX1 and AtRAX2, CaRAX2-like / SFL was renamed as CaRAX1/2a (Fig. **3d)**

### *CaRAX1/2a / SFL* acts in the I2 meristem controlling its activity

The *SFL* gene regulates the I2 meristem by repressing its activity, so that in the *sfl* mutants the I2 is active for longer, producing more flowers than in the wild-type. To understand where the *CaRAX1/2a / SFL* gene acts, we analysed the expression pattern of the *CaRAX1/2a / SFL* gene by *in situ* hybridization on vegetative and inflorescence apices of wild-type and *sfl-d* chickpea plants.

Inflorescence apices of the *sfl-d* mutant showed no signal when hybridized with a probe for the *CaRAX1/2a* gene (Fig. **4a**,**b**), as expected since is deleted in the mutant. When wild-type inflorescence apices were hybridized with *CaRAX1/2a*, signal was observed in I2 meristems as confirmed by including additional marker genes for I2 and for floral meristems in the experiments. Serial sections of inflorescences hybridized with probes for *CaRAX1/a2* or *CaVEG1*, (ortholog of *VEG1/MtFULc*, I2 meristem identity genes; Berbel et al., 2012; Cheng et al., 2018) exhibited an overlapping pattern, showing that *CaRAX1/2a* is expressed throughout the I2 meristem (Fig. **4a)**. Moreover, *VEG1* expression is not affected in the *sfl* mutant, suggesting that that *SFL* does not affect specification of I2 meristems.

In contrast, when serial sections of wild-type inflorescence apices were hybridized with probes for *CaRAX1/2a* or *CaPIM* (ortholog of *PIM/MtPIM*, floral meristem identity genes; Taylor et al., 2002; Benlloch et al., 2006) both probes exhibited a distinct pattern, with *CaPIM* hybridizing in the floral meristem and the *CaRAX1/2a* signal being found in the I2 meristem from which the floral meristem was emerging (Fig. **4b)**. While largely complementary, *CaRAX1/2a* and *CaPIM* expression was not mutually exclusive, since some overlap in the signal of *CaRAX1/2a* and *CaPIM* was found in the boundary between the I2 and the floral meristem (Fig. **4b)**.

In contrast to *RAX* genes from Arabidopsis and tomato (Muller et al., 2006; Bush et al, 2011), expression of *CaRAX1/2a* was not detected in vegetative axillary meristems or leaf axils neither at the inflorescence apex nor at the vegetative apex (Fig. **4a**,**c**).

These results show that *CaRAX1/2a* is specifically expressed in the I2 meristems, and not in the I1, floral or axillary meristems. This further supports that *CaRAX1/2a* corresponds to the *SFL* gene, which specifically regulates I2 meristem activity.

## DISCUSSION

Our results show that the *SFL* gene, whose mutation cause the formation of a larger number of flowers in the chickpea inflorescence, the double-pod phenotype, corresponds to *CaRAX1/2a*, which encodes a R2R3 MYB transcription factor. In the inflorescence apex, *CaRAX1/2a* is specifically expressed in the I2 meristem, in agreement with its role of regulating the activity of that meristem. Its expression largely overlaps with that of *CaVEG1*, whose homologs in other legumes specify I2 meristem identity (Berbel et al., 2012; Cheng et al., 2018). That in the *sfl-d* mutant, expression of *CaVEG1* is not affected, suggests that *CaVEG1* acts upstream of *CaRAX1/*2a, activating its expression. Also, that expression of *CaRAX1/2a* is not found in the I1 meristem, together with the fact that the *sfl* mutations seem to specifically affect I2 activity, indicate that that the genetic networks that the control meristem activity differs for I1 and I2.

In general, the expression pattern of *CaRAX1/2a* in shoot meristems has some similarities but also marked differences with that of *RAX1/3* (*REGULATORS OF AXILLARY MERISTEMS1 and 3*) and *Blind* genes, its homologs in Arabidopsis and tomato, respectively (Keller et al., 2006; Muller et al., 2006; Bush et al., 2011). The Arabidopsis and tomato genes are expressed in the axils of vegetative leaves. Though *CaRAX1/2a* is expressed in the I2 meristems, which form in the axils of inflorescence leaves, its expression is not detected in vegetative axillary meristems or in the axils of vegetative leaves. Moreover, while the Arabidopsis and tomato genes are expressed transiently, at the very beginning of axillary meristem initiation, expression of *CaRAX1/2a* is not transient and it is observed in the I2 meristem throughout its development, supporting its role in controlling its maintenance.

According to their expression pattern, the Arabidopsis *RAX1/2/3* and tomato *Blind* genes act as regulators of vegetative axillary meristems, promoting their initiation (Keller et al., 2006; Muller et al., 2006; Bush et al., 2011). The recently described *Marchantia polymorpha GCAM1* (*GEMMA CUP-ASSOCIATED MYB1*) gene, encodes a R2R3 MYB from subgroup 14, where RAX and Blind proteins are included (Dubos et al., 2010). When overexpressed, *GCAM1* generates abnormal *M. polymorpha* plants with cell clumps where apparently tissue differentiation is suppressed and proliferation of undifferentiated cells is induced (Yasuy et al., 2019). From these results, GCAM1 was proposed to promote the establishment and maintenance of cell groups with low differentiation level, but with competence to proliferate, somehow resembling the activity of the Arabidopsis and tomato RAX genes, which promote the initiation of meristems (Yasuy et al., 2019). The chickpea protein CaRAX1/2a also acts a meristem regulator but, rather than as a meristem promoter, it acts limiting the proliferative period and the activity of the I2 meristems, apparently in contrast with Arabidopsis and tomato RAX proteins. It is possible that the chickpea RAX protein interacts in a different way with one or more central regulatory factors of the genetic machinery for meristem functioning. Interestingly, when *M. polymorpha CGAM1* was expressed in the Arabidopsis *rax1 rax2 rax3* mutant under the control of the *RAX1* promoter, it did not promote meristem formation but further inhibited it. However, expression of a truncated CGAM1 protein notably recovered axillary meristem formation in the triple mutant (Yasuy et al., 2019). This shows that a change in the *M. polymorpha* RAX protein sequence turned it from a negative to a positive meristem regulator, and supports the hypothesis that the different, inhibiting activity of CaRAX1/2a on meristem regulation might be due to differences in its protein sequence.

What is the relevance of *CaRAX1/2a* function in the activity of the I2? What could be its contribution to control the number of flowers per I2 in nature? The *sfl* phenotype, an increase from one to two flowers, seems moderate. Besides, though *sfl-d* is a null mutation, the expressivity of the double-pod phenotype is not complete, since not every I2 in a *sfl* plant produces two flowers (Kumar et al., 2000). Arabidopsis *RAX1/2/3* genes have been shown to act redundantly in the promotion of axillary meristems (Muller et al., 2006). As chickpea *RAX* genes have duplicated, it is not unlikely that there is redundancy between *CaRAX1/2a* and other *CaRAX* genes in the regulation of I2 meristem activity. Moreover, another gene, *CYM*, has been described in chickpea that repress the production of flowers by the I2 (Srinivasan et al., 2006). It is thus likely that the combined action of *CaRAX1/2a* with *CYM* and other *CaRAX* genes jointly determines the number of flowers formed at the I2.

As in chickpea, genotypes with a specific increase of the number of flowers in the I2 are also found in other legume species (Murfet, 1985; Mishra et al., 2019), suggesting that the function of *CaRAX1/2a / SFL* could be conserved in other legumes. For instance, in pea, mutation in either *Fn* or *Fna* increase the number of flowers in the I2. Similar to *carax1/2a / sfl-d*, effect of the individual *fn* and *fna* mutations on floral production is modest but the phenotype is enhanced in the double *fn fna* mutant, which exhibits a multipod phenotype (White, 1917; Lamprecht, 1947). As the pea genome contains close homologues to *CaRAX1/2a*, it is tempting to speculate that *Fn* and *Fna* could be *RAX* genes. In addition, the fact that the function of other genes regulating floral and inflorescence development in legumes is generally very conserved (Benlloch et al., 2015; Cheng et al., 2018; Roque et al., 2018) also supports that the function of *CaRAX1/2a / SFL* might be conserved in other legume species.

The *SUPERMAN* (*SUP*) gene, encoding a C2H2 zinc-finger transcriptional repressor, restricts the proliferation of floral organs in Arabidopsis flowers (Hiratsu et al., 2002). It has been recently described that *MtSUPERMAN* (*Mtsup*), its *Medicago truncatula* orthologue, also regulates proliferation of floral organs but in addition regulates I2 meristem activity. Thus, *mtsup* plants produce an increased number of abnormal flowers in their I2s, which rather than forming a stub, get converted into a terminal flower (Rodas et al., 2020). Although it is not its only function, MtSUP, similarly to CaRAX1/2a, contributes to restrict I2 meristem activity, suggesting that both proteins might cooperate in that function.

Further analysis of legume *RAX* genes, whether they are functionally conserved among this family, or whether they act redundantly in the regulation of I2 meristem activity, promise to lead to valuable knowledge to design useful tools to improve seed yield but also to help understanding the basis for variety on number of flowers per I2 among different legume species to generate morphological diversity.

## ACKNOWLEDGMENTS

We thank Dr Ludovico Dreni for advice on phylogenetic analysis and MJ Domenech for efficient technical assistance. We also thank Dr C Ferrándiz and Dr MA Blázquez helpful discussions and critical reading. Work at F.M. lab was financed through grants from the Spanish Ministerio de Ciencia Innovación y Universidades and FEDER (BIO2015-64307-R and PGC2018-099232-B-I00). Work at IFAPA and UCO has been supported by INIA project RTA2017-00041, co-financed by the European Union through the ERDF2014–2020 “Programa Operativo de Crecimiento Inteligente”. C.C. acknowledges her Ph.D. fellowship INIA-CCAA.

## Supplementary figures

**Fig S1.**
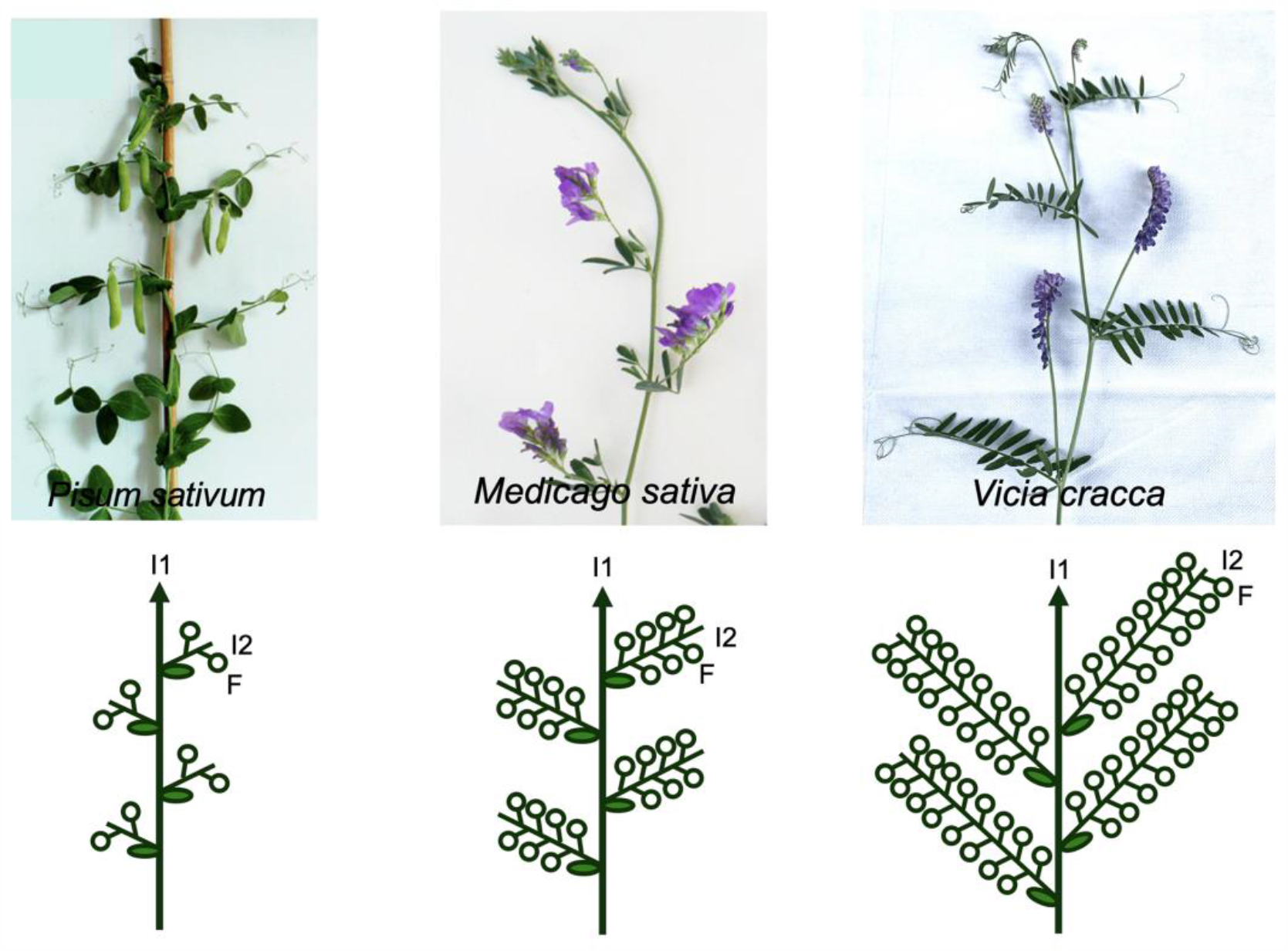
Legume species with different number of flowers in the I2. Inflorescences of legume species that produce a different number of flowers in their lateral I2s (top row). Diagrams of the inflorescences of those species (bottom row). I1, primary inflorescence; I2, secondary inflorescence; F, flower.

**Fig S2.**
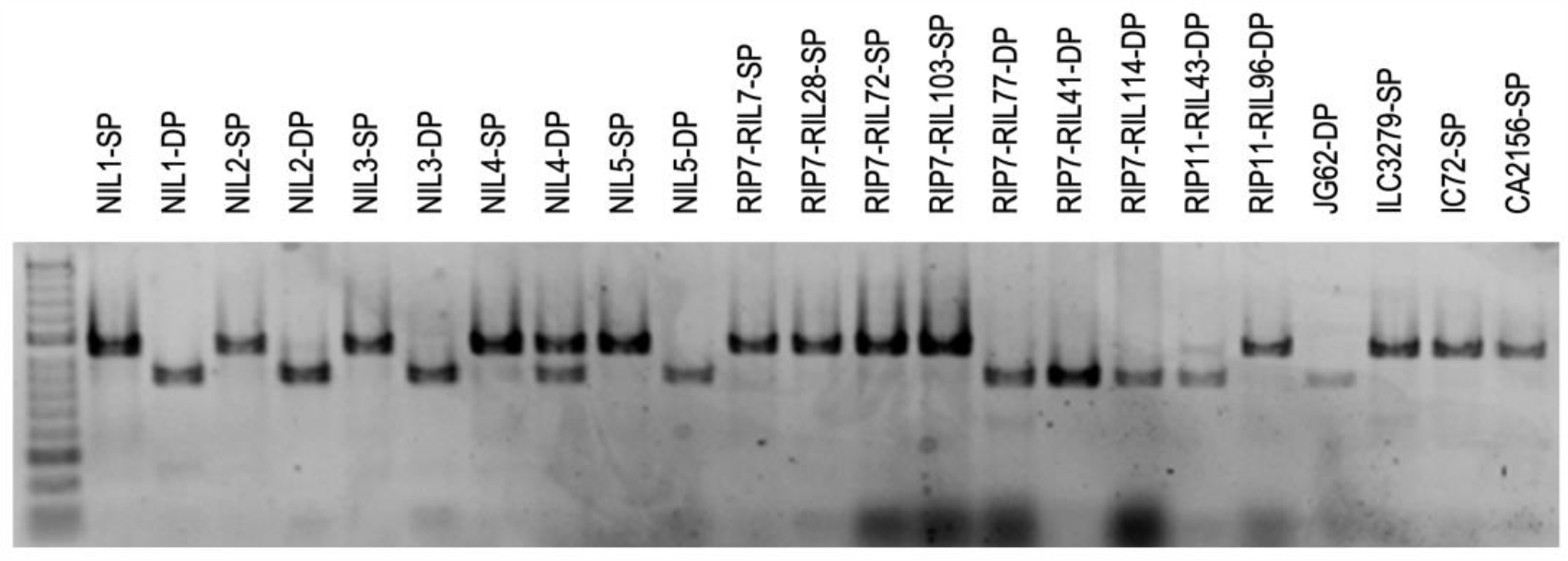
Amplification of the *CaRAX2-like* gene in single-pod and double-pod genotypes. PCR amplification with primers for the *CaRAX2-like* gene in diffferent chickpea lines that exhibit either a single-pod or a double-pod phenotype. SP, single pod; DP, double pod; NIL, nearly isogenic line; RIP, recombinant inbred population RIL, recombinant inbred line.

**Fig S3.**
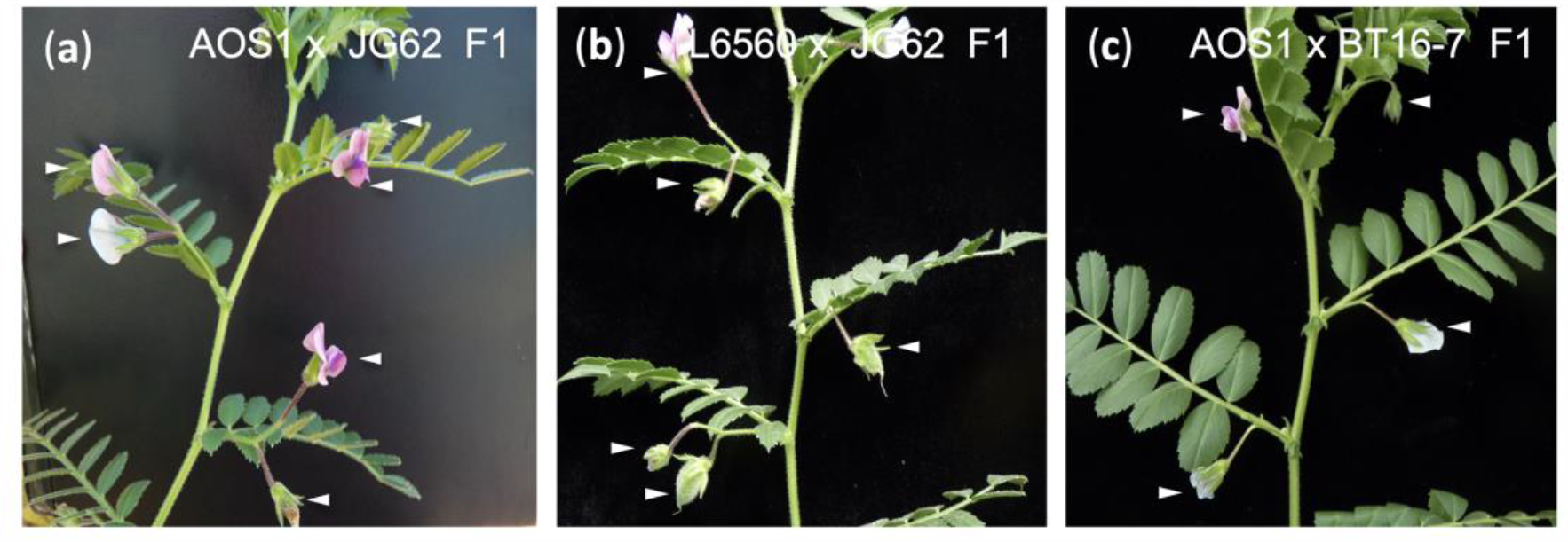
Allelic relationship between double-pod USDA mutants and the JG62 double-pod mutant. (a, b) Inflorescence phenotype of F1 plants derived from crosses of the USDA double-pod lines AOS1 (a) and L6569 (b) with the *sfl-d* double-pod parental line JG62. The F1 plants had I2s with two flowers (arrowheads). (c) F1 plant derived from a cross of the single-pod parental line BT16-7 with the USDA double-pod line AOS1. The F1 plants had I2s with a single flower.

## Supplementary tables

**Table S1.**
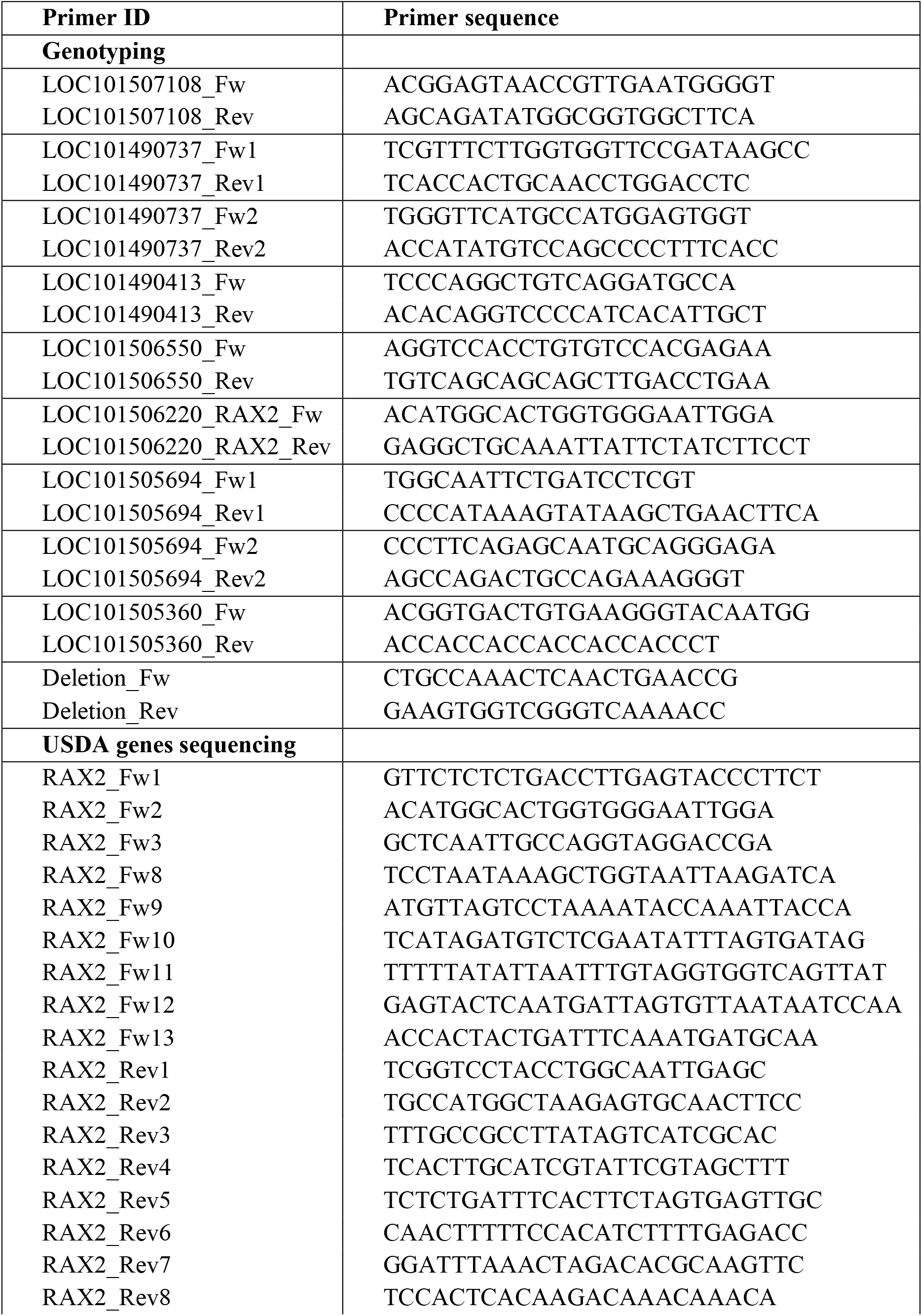
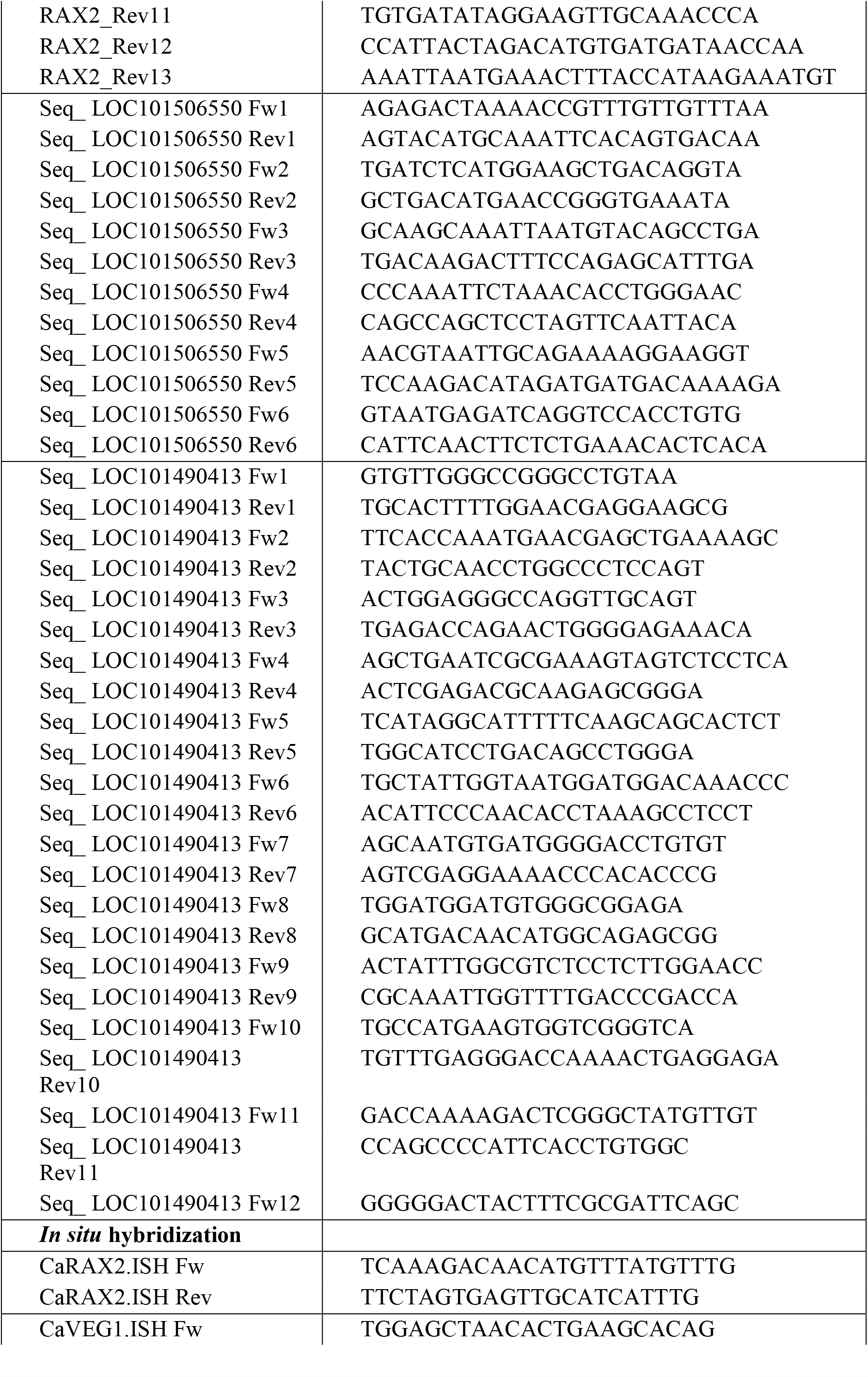
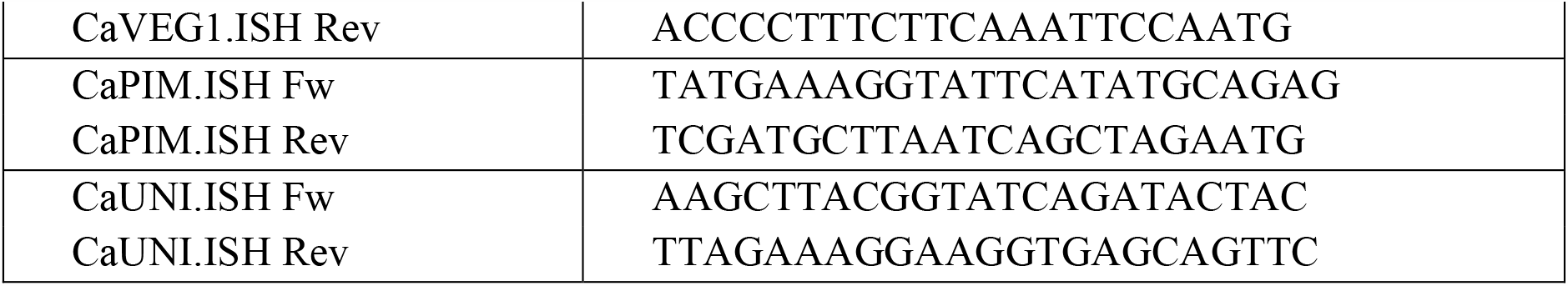
List of primers used in this study.

**Table S2.** Accession number of the genes in the phylogenetic tree in Fig 3d.

| **Species** | **Protein** | **Accession** |
| --- | --- | --- |
| *Arabidopsis thaliana* | AtRAX1<br>AtRAX2<br>AtRAX3<br>AtMYB35<br>AtMYB36<br>AtMYB68<br>AtMYB80<br>AtMYB87 | AtRAX1_AtMYB37_AT5G23000_S14<br>AtRAX2_AtMYB38_AT2G36890_S14<br>AtRAX3_AtMYB84_AT3G49690_S14<br>AtMYB35_AT3G28470_NS<br>AtMYB36_AT5G57620_S14<br>AtMYB68_AT5G65790_S14<br>AtMYB80_AT5G56110_NS<br>AtMYB87_AT4G37780_S14 |
| *Solanum lycopersicum* | SlyBLIND | SI_NP_001234643.1 BLIND_AAL69 |
| *Cicer arietinum* | CaRAX1/2a<br>CaRAX1/2b<br>CaRAX1/2c<br>CaRAX3a<br>CaRAX3b<br>CaMYB36 | Ca_XP_004506057.1<br>Ca_XP_004487066.1<br>Ca_XP_004494064.1<br>Ca_XP_004511803.1<br>Ca_XP_027189265.1<br>Ca_XP_004507824.1 |
| *Lotus japonicus* | LjRAX1/2c<br>LjRAX3a<br>LjRAX3b<br>LjMYB36 | Lj1g3v4593270.1<br>Lj2g3v1983980.1<br>Lj0g3v0236339.1<br>Lj4g3v2079370.1 |
| *Medicago truncatula* | MtRAX1/2a<br>MtRAX1/2b<br>MtRAX1/2c<br>MtRAX3a<br>MtRAX3b1<br>MtRAX3b2<br>MtMYB36 | Mt_XP_013455940.1<br>Mt_XP_013465364.1<br>Mt_XP_003625681.1<br>Mt_XP_003611533.1<br>Mt_XP_013461827.1<br>Mt_XP_024630749.1<br>Mt_XP_003610162.1 |
| *Pisum sativum* | PsaRAX1/2a<br>PsaRAX1/2b<br>PsaRAX3a<br>PsaRAX3b<br>PsaMYB36 | Psat7g146760.1<br>Psat5g280960.1<br>Psat2g172560.1<br>Psat0s127g0040.1<br>Psat4g012160.1 |

